# DNA methylation variability provides a complementary epigenetic signature of aging heterogeneity: Findings from the Canadian Longitudinal Study on Aging and the Baltimore Longitudinal Study of Aging

**DOI:** 10.1101/2025.08.25.671156

**Authors:** Olga Vishnyakova, Joosung Min, Ann Z. Moore, Toshiko Tanaka, Luigi Ferrucci, Xiaowei Song, Kenneth Rockwood, Angela Brooks-Wilson, Lloyd T. Elliott

## Abstract

**BACKGROUND:** Human aging does not follow a single trajectory. Epigenetic changes offer insight into the heterogeneity in aging by reflecting the combined influence of genetic, environmental, and lifestyle factors on the timing and progression of age-related changes beyond what chronological age alone can explain. Recent studies in cancer and aging underscore the importance of methylation variability as a marker of biological dysregulation. METHODS: We investigated the role of DNA methylation in aging heterogeneity by performing epigenome-wide differential methylation and variance association analyses in blood samples from 1,445 Canadians aged 45 to 85 from the Canadian Longitudinal Study on Aging.

**RESULTS:** We identified 448 differentially methylated regions and 488 differentially variable regions associated with health decline as measured by the health deficit accumulation Frailty Index, cognitive function, and physical function. These two classes of regions showed minimal overlap, with distinct gene coverage, suggesting that variability contributes a complementary signal to aging heterogeneity. Genes overlapped by differentially methylated regions were enriched for immune and inflammation-related pathways, whereas differentially variable regions highlighted additional localized, CpG island–enriched signals shared across health domains, consistent with regionally structured rather than diffuse dysregulation. By integrating significant CpGs from both analyses, we constructed an epigenetic biomarker. The biomarker was associated with all-cause mortality and showed higher discrimination than biomarkers constructed from differential methylation or variability alone, with a similar pattern reproduced in the Baltimore Longitudinal Study of Aging.

**CONCLUSIONS:** These findings suggest that DNA methylation variability may provide a complementary dimension of epigenetic aging and support further evaluation in larger cohorts with more mortality events.

## Introduction

Human aging follows no single trajectory. Longer life expectancy has led to a substantial rise in both the number and proportion of older adults, yet the burden of chronic disease remains high (1). At the same time, aging is highly heterogeneous, with wide inter individual differences in healthspan. Understanding the sources of this variation is essential for identifying mechanisms that influence functional decline and for extending the years lived free of disease and disability. Evidence from twin studies suggests that genetics accounts for approximately 25% of lifespan variation (2), indicating that environmental, behavioral, and social factors play a substantial and potentially modifiable role, in part through epigenetic mechanisms.

Epigenetic modifications regulate gene expression without altering the DNA sequence (3). These changes are influenced by aging, environment, lifestyle, and disease, and serve as a functional link between genetic and non-genetic factors. DNA methylation, one of the most studied epigenetic marks, affects DNA accessibility and transcriptional activity (4,5). Age-associated shifts in DNA methylation patterns have consistently been observed across tissues (6,7). The fact that individuals of the same chronological age, including monozygotic twins, show substantial differences in methylation profiles supports the view that DNA methylation marks molecular variation underlying aging heterogeneity (8).

DNA methylation biomarkers capture aggregated molecular changes associated with aging. Early models identified CpG (cytosines preceding guanines in DNA) sites whose methylation levels correlate with chronological age (6,9,10), while later approaches included sites linked to health status, disease risk, and physiological markers such as blood chemistry, systolic blood pressure, and lung function (11–13). These biomarkers have shown strong associations with mortality and age-related diseases (e.g., cardiovascular disease, type 2 diabetes, cancer, Alzheimer’s disease) and respond to behavioral and clinical interventions (14,15). Despite varying methodologies, the focus remains on differential methylation at specific sites, potentially overlooking other methylation dynamics.

DNA methylation in normal tissues tends to be relatively stable, whereas cancer tissues often display markedly increased variability, suggesting that altered variance can reflect disrupted regulatory states (16). Changes in variability appear also to inform complex phenotypes beyond cancer. In our previous work, we found that variance heterogeneity between least healthy and most healthy older adults can reveal traits under homeostatic regulation, which we proposed to be closely related to overall health and healthy aging (7,17,18). These observations indicate that methylation variability may capture biological processes relevant to aging heterogeneity and motivate examining variability as a complementary dimension of the methylome.

The extent to which DNA methylation variability contributes to aging heterogeneity remains unclear. To address this question, we performed an epigenome wide variance association analysis in blood samples from 1,445 Canadians aged 45 to 86 to identify CpG sites with differential methylation variability. We also conducted an epigenome wide association study of methylation levels and health decline, and examined gene and pathway level enrichment to contextualize CpGs identified in both analyses. Finally, we integrated significant CpGs from the two analyses to construct a DNA methylation biomarker and evaluated its performance in relation to healthspan, including replication in an independent cohort from the Baltimore Longitudinal Study of Aging.

## Methods

### Data

We examine data from the Canadian Longitudinal Study on Aging (CLSA) (19). These data involve a stratified sample of 51,338 Canadian males and females aged 45 to 85 years. Community-dwelling participants were recruited by CLSA from the ten Canadian provinces, excluding individuals unable to respond in English or French; residents of the three Canadian territories and some remote regions; individuals living on First Nation reserves and other First Nations settlements in the provinces or in nursing homes; full-time members of the armed forces; and individuals with significant cognitive impairment at recruitment. The subsample of participants selected for the Comprehensive Cohort of the CLSA underwent detailed physical assessments and provided blood and urine samples. Baseline data were collected between May 2012 and July 2015. The average follow-up period was six years, defined as the time between baseline and Follow-up 2 data collection. The onset of age-related diseases after baseline was determined using data from both Follow-up 1 and Follow-up 2 assessments. All participants of CLSA provided informed consent prior to data collection. The present study was approved by the joint Research Ethics Board of the BC Cancer and the University of British Columbia, and harmonized with Simon Fraser University (#H22-01012).

Genome-wide DNA methylation profiling was performed by CLSA on peripheral blood mononuclear cells from 1,478 participants who provided blood samples at baseline, using Illumina Infinium MethylationEPIC BeadChip arrays. Raw signal intensities underwent rigorous preprocessing, including sample- and probe-level quality control, outlier detection, and exclusion of poorly performing probes, resulting in 1,445 high-quality samples and 783,136 probes (20). Immune cell proportions were estimated using a reference-based deconvolution algorithm (21) and used to adjust methylation values. We examined remaining samples for relatedness and mismatch between self-reported and genetic sex. Of the total sample, 1,284 (89%) participants were classified as having European ancestry, according to principal component analysis of genetic data conducted by CLSA (22). We removed CpG sites with a SNP within 10 bp of the probe as such variants can bias methylation estimates in array-based assays (23). Characteristics of the CLSA study sample at baseline are provided in Table S1.

### Replication cohort

Replication analyses were conducted in the Baltimore Longitudinal Study of Aging (BLSA), a continuously enrolled cohort study of normative human aging initiated in 1958 (24). The BLSA protocol (Protocol number 03-AG-0325) was approved by the National Institutes of Health Intramural Research Program Institutional Review Board and all participants provide written informed consent at each study visit. The replication sample consisted of 728 individuals with Illumina 450k DNA methylation data and GWAS based ancestry information, drawn from visits occurring between April 2003 and March 2010. These visits served as the index visits for linkage to clinical outcomes and for defining follow up in survival analyses. Details on sample selection and phenotype definitions are provided in Table S2 and Supplementary Note 1.

### Statistics and Reproducibility

The study workflow is detailed in Figure S1. Primary analyses focused on the largest ancestral subset (European ancestry), representing 89% of the cohort. Health scores were adjusted for age. To perform inference and evaluate the performance of the epigenetic biomarkers, we randomly partitioned the participants with European ancestry into a training set (85%) and a validation (15%) set, resulting in sample sizes of 1,091 and 193, respectively. Stratification was based on sex and age group, producing balanced age and sex distributions between sets (Table S3). For all analyses, we used cell type adjusted methylation values. Estimated immune cell proportions were regressed from each CpG and the residuals were carried forward into all differential methylation, variance association, and biomarker analyses.

### Health status assessment

We assessed participants’ health status by measuring the deficit accumulation Frailty Index (25), cognitive function, and physical function. Because the cognitive and physical function tests differ in scale, distribution, and direction of risk, rank normalized scores were used to construct the composite instruments. The cognitive function was defined as the mean of three domain scores (memory, executive function, psychomotor speed), each derived from rank normalized domain specific tests. Physical function was defined as the mean of rank normalized scores from five assessment tests. To ensure comparability among these measures, all instruments were standardized to a unit interval, with higher values indicating greater health deficits. Details on phenotype definitions and scoring procedures are provided in Table S4. The construction and methodologies used for these instruments are described elsewhere (17).

### Differential methylation analysis

To investigate methylation patterns associated with aging heterogeneity, differential methylation analyses were conducted separately for each health score: Frailty Index, cognitive function, and physical function. M-values (logit-transformed beta-values) were used to stabilize variance across the methylation range. Differentially methylated positions (DMPs) are individual CpG sites where mean methylation levels are associated with health status. DMPs were identified using the *limma* package, which fits a linear model to each CpG site and applies empirical Bayes shrinkage to improve variance estimation (26). For each analysis, the model included the corresponding health score as the primary predictor, adjusting for age, sex, and the first three genetic principal components. Resulting p-values were adjusted for multiple testing using the Benjamini-Hochberg procedure to control the false discovery rate (FDR).

We identified differentially methylated regions (DMRs), genomic segments defined by aggregating multiple DMPs in close proximity that show coordinated associations with health decline, using *DMRcate*, which applies Gaussian smoothing (bandwidth = 1000 bp) to test statistics to account for spatial correlation across adjacent probes (27). Regions with a smoothed FDR < 0.05 were considered significant. Genes overlapping with DMRs were annotated to facilitate interpretation of biological pathways relevant to aging. Gene set enrichment analysis was performed using the *missMethyl* R package, which accounts for bias arising from the varying number of CpGs per gene on the Illumina MethylationEPIC v1 array (28).

### Variance association analysis

Differentially variable positions (DVPs) are individual CpG sites where methylation variance, rather than mean methylation level, is associated with health status. To identify DVPs, we conducted a variance association analysis. While prior studies compared methylation variance between discrete groups (8,29), we used double generalized linear models (DGLM) to identify DVPs for continuous health scores (30) implemented in the *dglm* R package (30). A DGLM jointly models both the mean and the variance of a response variable, allowing the variance to depend on covariates. Specifically, for a continuous outcome , i.e., DNA methylation of a particular CpG, the DGLM assumes:

Mean model: 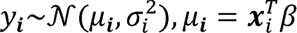

Dispersion model: 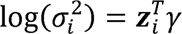

Here, **x***_i_*and **z***_i_* are covariate vectors for the mean and dispersion models, respectively. For each CpG site, methylation levels were expressed as M-values and modeled assuming Gaussian errors. The mean model was specified as:

M-value ∼ health score + sex + age + PC_1_ + PC_2_ + PC_3_,

and the dispersion model as:

log(variance) ∼ health score + sex + age + PC_1_ + PC_2_ + PC_3_.

Here the health score is one of the following continuous health scores used in the analysis: Frailty Index, cognitive function, or physical function, with higher values indicating worse health. The dispersion coefficient estimates the association between the health score and the log-variance of methylation at each CpG. Positive coefficients indicate increased variability in methylation with worsening health. Statistical significance was evaluated using two-sided z-tests. CpGs with positive dispersion coefficients and Benjamini-Hochberg adjusted *p*-values *<* 0.05 were classified as DVPs. Bias and inflation were controlled using a Bayesian method (31).

Differentially variable regions (DVRs) are genomic segments defined by clusters of nearby DVPs that exhibit coordinated changes in methylation variability associated with health status. DVRs were identified by aggregating significant positions located within 1 kb of each other using a spatial smoothing algorithm, in a manner similar to the differential methylation analysis.

### Epigenetic biomarkers

We developed a DNA methylation–based biomarker of aging to evaluate whether incorporating DVPs alongside DMPs improves the prognostic performance of methylation measures in relation to mortality and age-related diseases.

The biomarker was constructed in two steps, using separate training and test sets. In the first step, for each of three health scores—Frailty Index, cognitive function, and physical function—we fitted an elastic net regression model (32) to select CpGs and estimate methylation-based scores. We trained the following models using the *glmnet* R package with 10-fold cross-validation to select the regularization parameter λ (33):

health score ∼ DMPs + DVPs + age + sex,

with the output being predicted health score values per individual, representing a methylation-based health score. Each score was rescaled to match the mean and standard deviation of chronological age and then linearly regressed on chronological age to obtain residuals.

In the second step, we estimated weights for the three methylation-based scores using Cox proportional hazards regression:

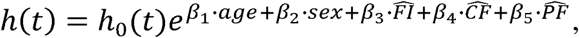

where, 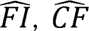, and 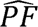 are the methylation-based Frailty Index, cognitive, and physical function scores from step one, h_0_ (*t*) is the baseline hazard function—representing the hazard when all covariates are set to zero. The coefficients β_1_, … ,β_5_ are the log hazard ratios associated with each covariate. The weighted sum of these components formed the final epigenetic age measure. This score was again regressed on chronological age, and the residual—epigenetic age deviation—served as the final biomarker.

To evaluate the prognostic value of the epigenetic biomarker, we assessed model discrimination in the test set using Cox models adjusted for age and sex. Prognostic performance was quantified using the concordance index (C-index).

To assess the contribution of CpGs derived from differential methylation and differential variable analyses, we constructed control models using only DMPs or only DVPs as inputs to the elastic net models in step one, following the same procedure. These models enabled evaluation of whether combining DMPs and DVPs improved discriminatory power. Additionally, we included a null model:

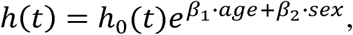

to test whether the biomarker added prognostic information beyond basic demographic covariates.

We further compared the biomarker against established benchmarks: Frailty Index; Hannum (10) and Horvath (9) clocks, included in the CLSA COM dataset; PhenoAge (34) and GrimAge v2 (11), computed using the *Biolearn* Python library (35); the methylation risk score (MRS), Shannon entropy, and the epigenetic frailty risk score (eFRS). A MRS was constructed by weighting CpGs with effect sizes derived from the differential methylated analysis. eFRSs were computed using the published pretrained model based on 20 preselected CpG predictors (36). Finally, we included a methylation Shannon entropy measure calculated over DVPs, as an index of epigenetic disorder, previously shown to increase with age and reflect reduced information content in the methylome (10).

For external validation in BLSA, we also constructed 450K-restricted versions of the DMP-DVP, DMP-only, and DVP-only biomarkers using the same CLSA training procedure but limiting predictors to CpGs available on the Illumina HumanMethylation450 array. These 450K-restricted models were used only for cross-platform validation, whereas the primary CLSA analyses used the full set of selected CpGs available in the CLSA methylation data.

### Heritability estimation

Genotyping was done by CLSA using the Affymetrix UK Biobank Axiom array, with phasing and imputation using TOPMed reference panel v.r2 (37) with marker-based quality control performed by CLSA (22). Variants were examined for discordant genotype frequency between batches, departure from HWE, discordance across control replicates, and sex genotype frequency discordance. We removed low-quality imputed genetic variants: with a minor allele frequency lower than 0.1%, imputation quality score<0.5 and missing rate>0.1. Remaining 11.7 million variants were used for genome-wide association study (GWAS). We performed a GWAS on epigenetic age deviation, using SAIGE (Scalable and Accurate Implementation of Generalized mixed model) (38) in a cohort of 1,284 unrelated individuals of European ancestry. The GWAS summary statistics were used as input for LD Score Regression (LDSC, version 1.0.1) (39) to estimate SNP-based heritability. LDSC was performed using European reference panel LD scores and regression weights, retaining 819,098 SNPs after merging. A two-step estimator with a cutoff of 30 was applied to handle outliers in test statistics.

## Results

### Health assessment and participants’ characteristics

We analyzed DNA methylation data from 1,284 participants aged 45 to 85 years (mean age 64 *±* 10 years, 51% female) enrolled in CLSA. Samples were collected by CLSA during the period from May, 2012 to July, 2015, with a mean tracking period of 6 years. Descriptive statistics and intercorrelations among health scores are presented in Figure 1A-B.

**Figure 1:**
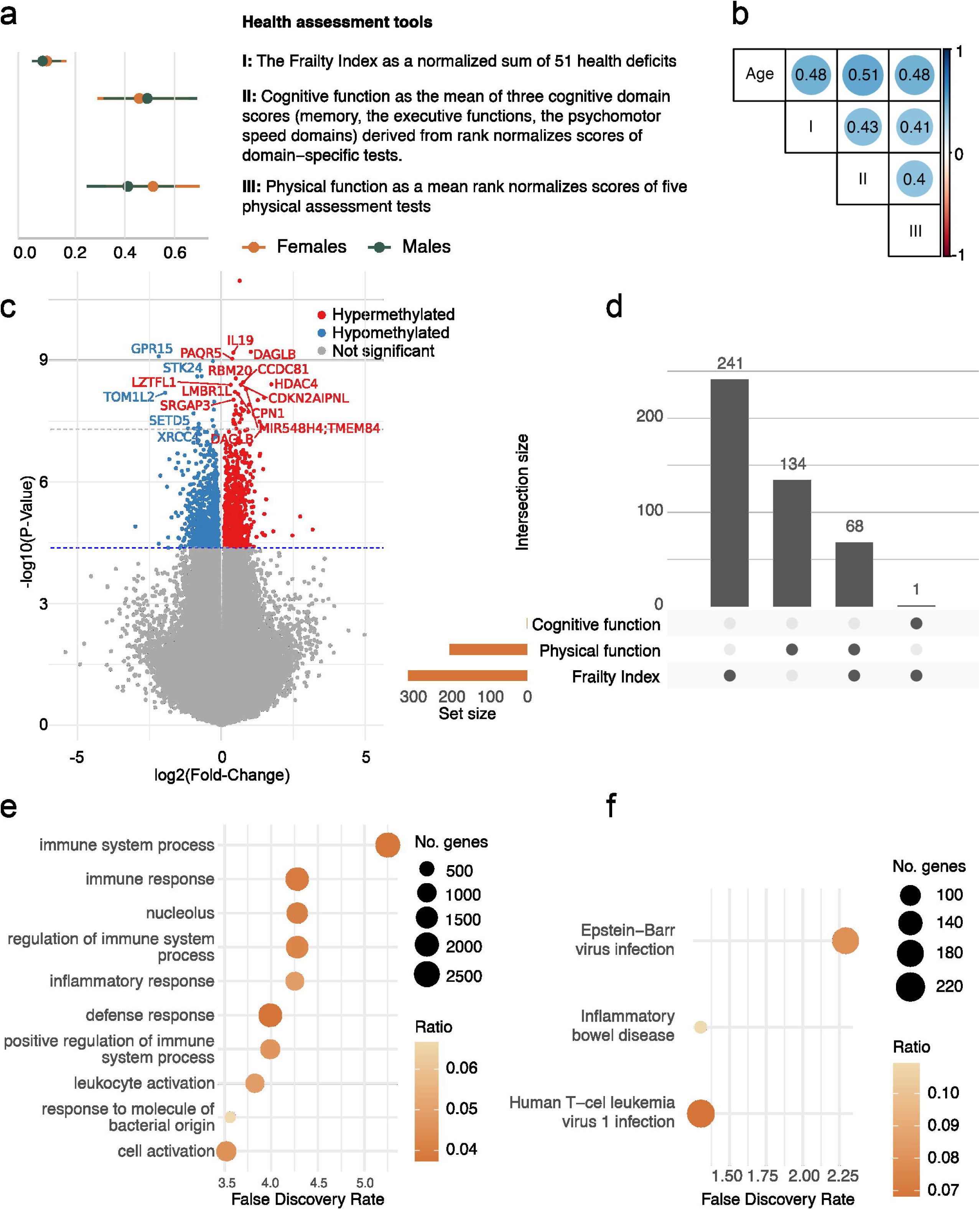
Differential methylation analysis results. (a) Mean and standard deviations of three health instruments, representing health status in different ways with scores ranging from 0 (healthiest) to 1 (least healthy); (b) Pearson correlations between health instruments and chronological age. All health deficit scores were strongly correlated with chronological age; (c) Volcano plot showing statistical significance (*−* log_10_ p-value) and effect size of all CpGs included in the differential methylation analysis; (d) The overlap of genes covered by differentially methylated regions (DMRs) associated with the Frailty Index, cognitive and physical function; (e) Top 10 significant gene ontology terms from gene set enrichment analysis of differentially methylated genes, with FDR-adjusted p-values from a one-sided Wallenius’ noncentral hypergeometric test; (f) Significant KEGG pathways from gene set enrichment analysis of differentially methylated genes, with FDR-adjusted p-values from a one-sided Wallenius’ noncentral hypergeometric test and dot size indicating the number of genes involved. Alt text: Six-panel summary of differential methylation analyses. The three health-deficit measures are intercorrelated and strongly correlated with chronological age. The analyses identified 1775 differentially methylated positions that passed FDR correction. Significant positions include both hypermethylated and hypomethylated CpGs. Most genes covered by differentially methylated regions are specific to the Frailty Index, with some overlap among the Frailty Index, cognitive function, and physical function. Enriched biological terms and pathways predominantly involve immune and inflammatory processes.

### Differential methylation regions highlight immune and inflammatory processes in aging heterogeneity

After applying quality control filters, 764,839 sites were tested for association with health scores in 1,091 individuals from the training set to investigate methylation patterns associated with aging-related health variation. We identified 1,775 differentially methylated positions (DMPs) that exceeded FDR correction, of which 70 surpassed the epigenome-wide significance threshold (40), 9.42 *×* 10*^−^*^8^ (Figure S2). Among these 70 loci, 26 positions were hypomethylated and 44 were hypermethylated (Figure 1C and Table S5) 52 of the 70 DMPs were mapped to gene regions, including the gene bodies of *DAGLB*, involved in lipid metabolism (cg17890233, *p* = 6.20 *×* 10*^−^*^10^), *IL19*, involved in immune signaling (cg16513119, *p* = 6.53 *×* 10*^−^*^10^), and *GPR15*, involved in immune cell trafficking (cg19859270, p = 8.09 *×* 10*^−^*^10^).

Beyond individual DMPs, we examined differentially methylated regions (DMRs), which span multiple contiguous DMPs. Because neighboring CpG sites are highly correlated, DMRs may offer greater biological relevance and stronger links to gene expression differences compared to isolated DMPs (29). We identified 484 DMRs (Figure 2 and Table S6). For the Frailty Index, the most significant region was chr19:47287778–47289611 (1834 bp, 12 CpGs, *p* = 9.68 *×* 10*^−^*^24^) overlapping *SLC1A5*, followed by chr6:30852963– 30854551 (1589 bp, 17 CpGs, *p* = 1.47 *×* 10*^−^*^16^) which contains the gene *DDR1*. Cognitive function showed a top region on chr11:2890551–2890725 (175 bp, 14 CpGs, *p* = 3.28 *×* 10*^−^*^14^) with no annotated gene, and a second region on chr22:45608686–45608713 (28 bp, 3 CpGs, *p* = 5.16 *×* 10*^−^*^14^) overlapping the gene *KIAA0930*. Physical function identified a highly significant region on chr19:47287778–47289611 (1834 bp, 12 CpGs, *p* = 2.97 *×* 10*^−^*^21^), also overlapping *SLC1A5*, followed by chr6:31507433–31508923 (1491 bp, 14 CpGs, *p* = 4.83 *×* 10*^−^*^18^) overlapping *DDX39B*. To compare DMR patterns across health measures, we examined overlap among genes covered by DMRs. Most mapped genes were unique to the Frailty Index (n = 241); 68 were shared with physical function, and *KIAA0930* was common to both Frailty Index and cognitive function (Figure 1D).

**Figure 2:**
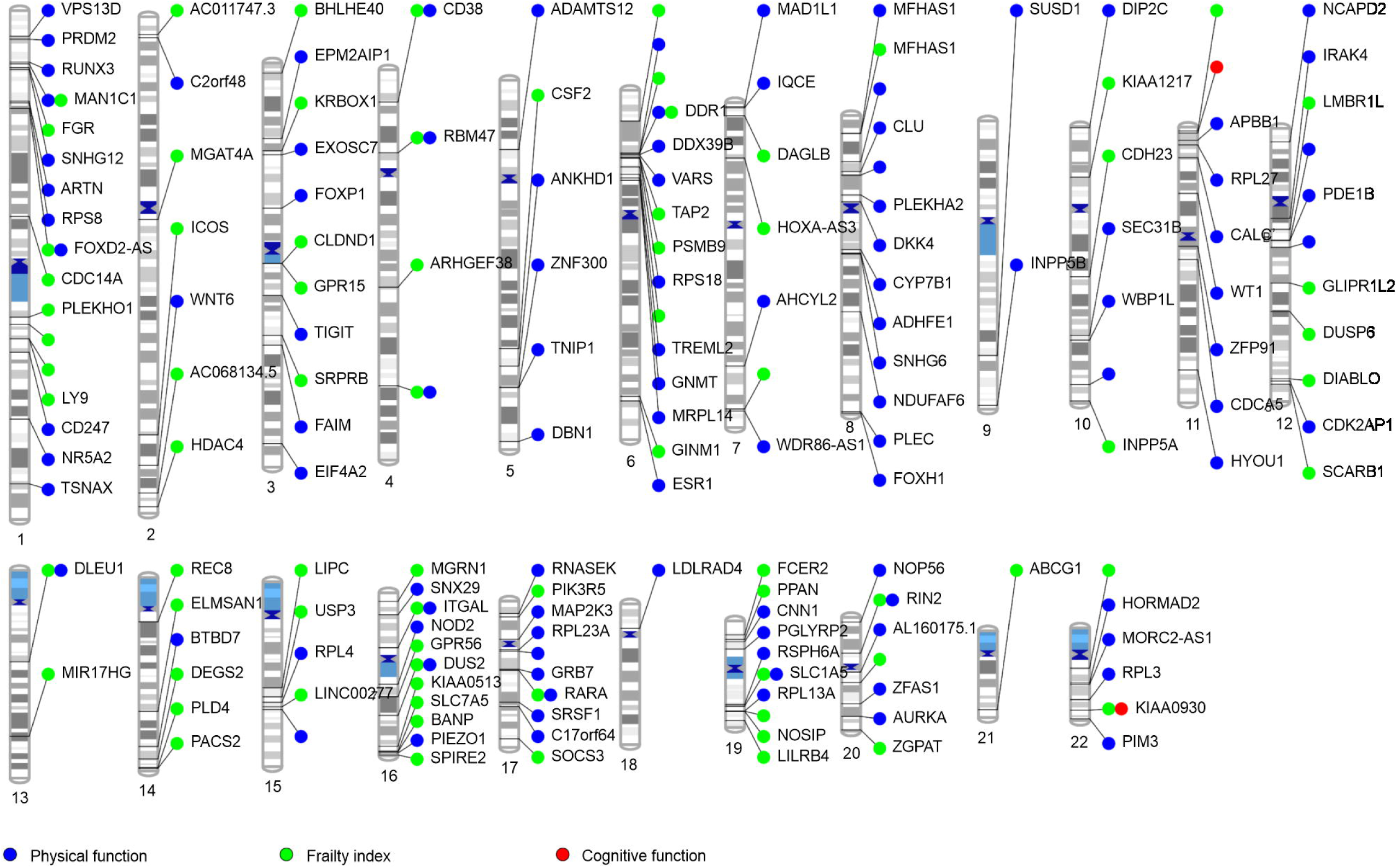
Chromosomal ideogram of differentially methylated regions associated with the Frailty Index, cognitive function and physical function (only regions with FDR<9.42 *×* 10*^−^*^8^ are shown). The plot was generated using PhenoGram (https://ritchielab.org/software/phenogram-downloads). Alt text: Chromosomal ideogram showing the genome-wide locations of differentially methylated regions associated with the Frailty Index, cognitive function, and physical function. Significant regions are distributed across multiple chromosomes. Prominent regions include *SLC1A5* for the Frailty Index and physical function and *KIAA0930* for cognitive function.

To gain biological insight into the functional relevance of DMRs, we performed gene set enrichment analysis. A total of 97 gene ontology (GO) terms were significantly enriched (Table S7). The top GO terms were related to immune and inflammatory processes, including immune system process (FDR = 5.44 *×* 10*^−^*^6^), inflammatory response (FDR = 5.26*×* 10*^−^*^5^), and regulation of immune system process (FDR = 9.38 *×* 10*^−^*^5^). We also conducted gene set enrichment analysis for KEGG pathways to further elucidate the biological roles of differentially methylated genes (Table S8). Three KEGG pathways were significantly enriched (FDR < 0.05), with the top pathway being Epstein-Barr virus infection (FDR = 0.005), followed by human T-cell leukemia virus 1 infection (FDR = 0.05). DMR-covered genes were enriched for human aging genes (41), with 14 of 305 human aging genes represented among the DMR-associated genes (*p* = 0.013). Results for the gene set enrichment analysis are shown in Figure 1E-F.

### Variance association analysis identifies localized regions shared across health domains

To investigate whether aging-related health decline is accompanied by altered methylation variability, we modeled CpG-specific methylation variance as a function of the Frailty Index, physical function, and cognitive function. We detected 473 DVPs surpassing the epigenome-wide significance threshold. At a less conservative FDR-corrected threshold, we identified 3,197 DVPs (Figure S3 and Table S9).

Next, we aggregated adjacent CpGs, yielding 673 differentially variable regions overlapping 591 unique genes across the genome (Figure 3 and Table S10). In contrast to DMRs, DVR-covered genes showed substantial overlap between the different health measures, with *CCDC171*, *VILL*, and *ZNF783* shared across all health scores (Figure 4A). To explore biological relevance, we performed gene set enrichment analysis; however, no GO terms or KEGG pathways remained significantly enriched after FDR correction.

**Figure 3:**
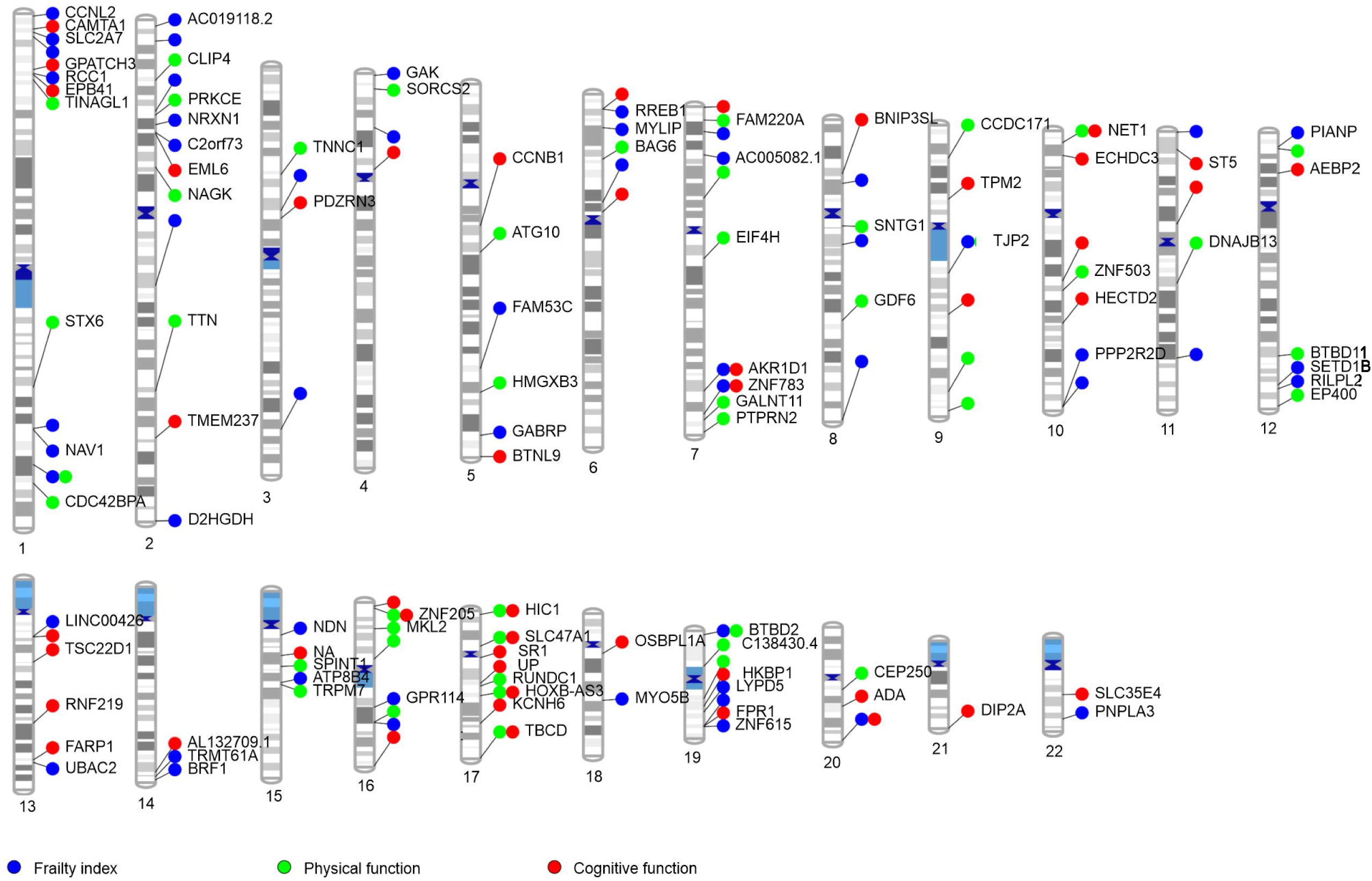
Chromosomal ideogram of differentially variable regions associated with the Frailty Index, cognitive function and physical function (FDR<9.42 *×* 10*^−^*^8^). Alt text: Chromosomal ideogram showing the genome-wide locations of differentially variable regions associated with the Frailty Index, cognitive function, and physical function. Significant regions are distributed across the genome and include regions shared among the three health measures.

**Figure 4:**
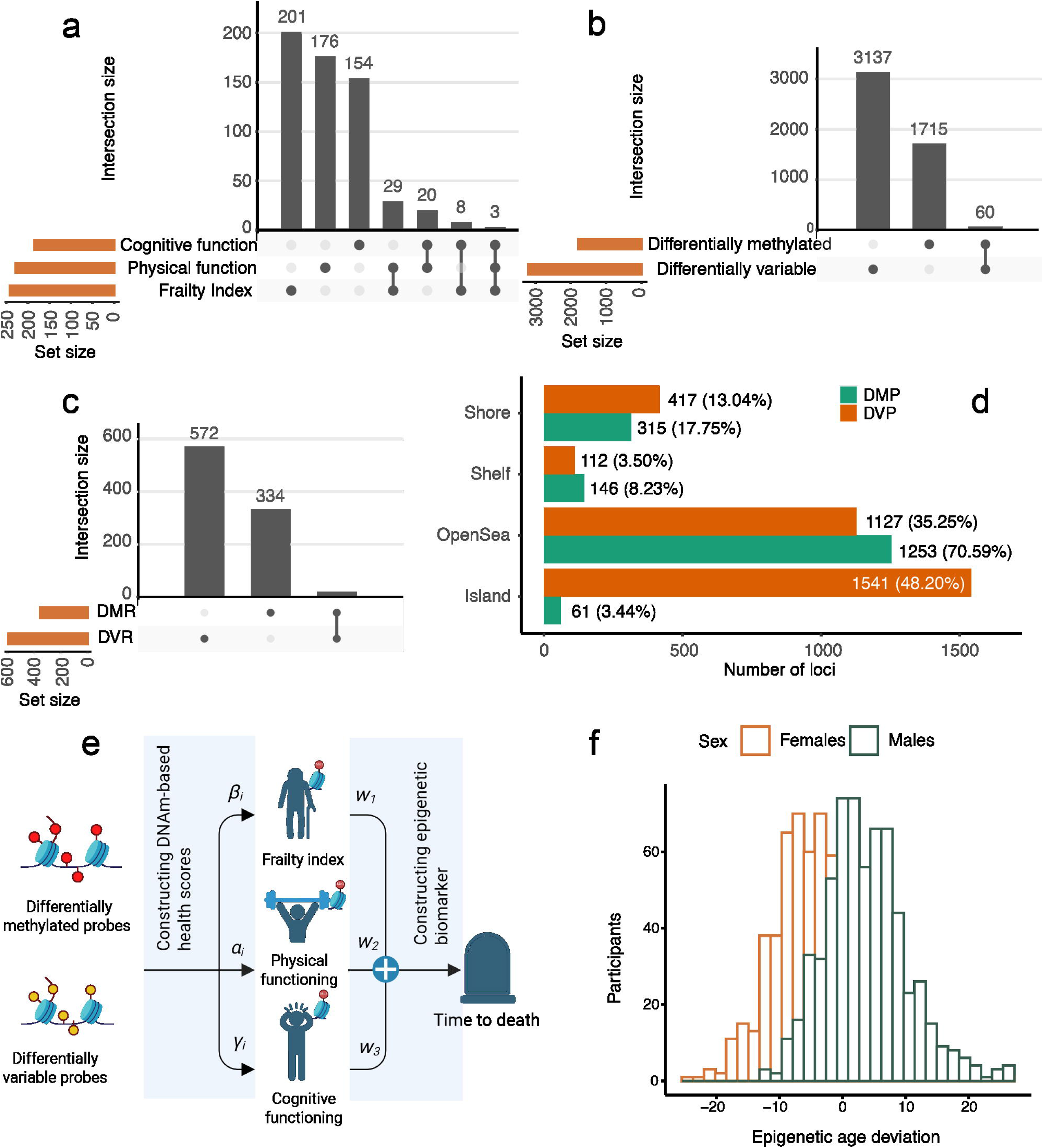
DNA methylation variability captures unique aging signals. (a) The overlap of genes covered by differentially variable regions (DVRs) associated with the Frailty Index, cognitive and physical function; (b) Overlap between differentially variable positions (DVPs) and differentially methylated positions (DMPs); (c) Differences in the genomic regions associated with differential methylation and variability. Overlap between genes within differentially variable regions (DVRs) and differentially methylated regions (DMRs); (d) Location of DVPs and DMPs relative to their proximity to CpG islands; (e) DVP-DMP-based epigenetic biomarker approach overview; (f) Epigenetic age deviation sex-specific histogram. Alt text: Six-panel comparison of differential methylation and methylation variability signals. Differentially variable regions show greater gene overlap across the three health measures than differentially methylated regions. Differentially variable positions are enriched in CpG islands and promoter-proximal regions, whereas differentially methylated positions are enriched in open-sea regions and gene bodies. The final panels illustrate construction of the combined DMP-DVP epigenetic biomarker and the sex-specific distribution of epigenetic age deviation.

### Differentially variable regions are distinct from differentially methylated regions

Differential methylation and genome-wide variance analyses revealed distinct epigenetic patterns through DMPs/DVPs and DMRs/DVRs (Figure 4B-C). Genomic annotation enrichment analyses showed distinct genomic localization patterns for differential methylation and variability signals (Table S11, Figure 4D and Figure S4). DVPs were strongly enriched in CpG islands relative to the tested CpG background (OR = 4.19, FDR < 10*^−^*^130^), whereas DMPs were enriched in open sea regions (OR = 1.84, FDR = 1.43 × 10 ³³). Functional genomic region enrichment further showed that DVPs were preferentially localized to promoter-proximal and first-exon regions, including TSS200 (OR = 2.57, FDR = 3.38 × 10 ¹ ¹), 1stExon (OR = 2.54, FDR = 2.51 × 10 ), 5′UTR (OR = 1.44, FDR = 7.65 × 10 ² ), and TSS1500 (OR = 1.08, FDR = 0.034), whereas DMPs were enriched only in gene bodies (OR = 1.67, FDR = 2.33 × 10 ³ ).

The two classes of regions also differed in gene coverage: 334 DMR-only genes, 572 DVR-only genes, and 19 shared genes, including *HDAC4* and *TREML2*. DMRs spanned more CpGs (mean 5.4, SD 3.7) and wider ranges (mean 518 bp, SD 394) than DVRs (mean 4 CpGs, SD 3.0; mean 365.7 bp, SD 308.3). Expression quantitative trait methylation (eQTM) analysis (42) identified 32 DMPs associated with expression of 33 genes and 61 DVPs associated with expression of 94 genes (*p* < 9.42 *×* 10*^−^*^8^). The DMP- and DVP-eQTM annotations were largely distinct, sharing only *ARHGAP35* (Table S12 and Table S13).

### Non-linear analysis did not identify DNA methylation sweet spots

To explore potential non-linear relationships between DNA methylation and health decline, we applied segmented regression with a single breakpoint to DNA methylation levels across all CpG sites. This approach aimed to identify “sweet spots”—methylation levels associated with optimal health—where deviations in either direction link to worse health (17). However, none of the breakpoint models remained significant after FDR correction, suggesting that methylation variation associated with aging heterogeneity is primarily linear rather than reflecting an optimal value.

### An epigenetic biomarker integrating DMPs and DVPs captures complementary mortality-related signal

To evaluate whether the distinct methylation signals identified by differential methylation and variance association analyses could be summarized at the individual level, we constructed an epigenetic biomarker integrating CpGs selected from both analyses (Figure 4E-F). Briefly, selected CpGs were first used to derive methylation-based health scores for the Frailty Index, cognitive function, and physical function, which were then combined using Cox proportional hazards regression with all-cause mortality as the outcome. Training-set performance for the unrestricted and 450K-restricted elastic-net models used to derive methylation-based health scores is summarized in Table S14. The final DMP-DVP biomarker retained 789 CpGs after elastic-net selection (comprising 313 DMPs and 488 DVPs, with 12 CpGs shared by both classes), with limited CpG-level overlap with established epigenetic clocks, consistent with a distinct feature set derived from differential methylation and variability analyses (Table S15; Supplementary Note 2). Strong correlations with health instruments were observed in the training set, with weaker but significant correlations in the validation set (Table S16). Pairwise correlations between predictors in both the training and validation sets are shown in Figure S5.

In the held-out CLSA validation set, the integrated DMP-DVP biomarker was associated with all-cause mortality over six years (N = 187; 14 deaths; HR = 1.125, 95% CI: 1.015–1.247, *p* = 0.024; C-index = 0.877). Biomarkers constructed from DMPs alone or DVPs alone were also associated with mortality but showed lower discrimination in this sample. Benchmark epigenetic models were not significantly associated with mortality in the CLSA validation set (Table 1).

**Table 1:**
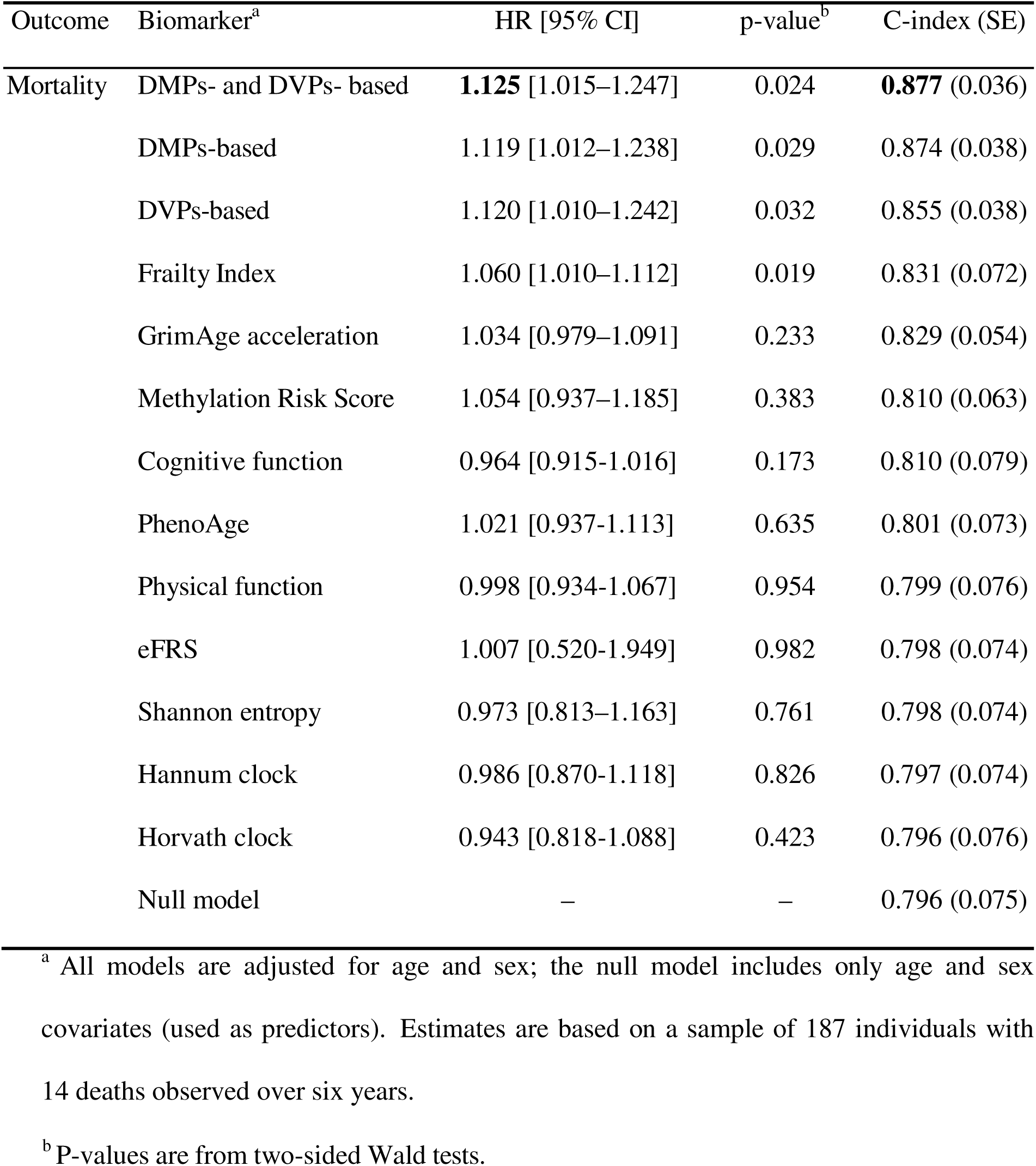
Hazard Ratios (HR) and concordance indices (C-index) for six year all cause mortality across aging biomarkers. The model based on both DMPs and DVPs has the most predictive hazard ratio and C-index. All models considered have a more predictive C-index than the null model. The model based on both DMPs and DVPs outperforms the models based on DMPs or DVPs alone, indicating distinct signal in these two modalities.

| Outcome | Biomarker<sup>a</sup> | HR [95% CI] | p-value<sup>b</sup> | C-index (SE) |
| --- | --- | --- | --- | --- |
| Mortality | DMPs- and DVPs- based | **1.125** [1.015–1.247] | 0.024 | **0.877** (0.036) |
|  | DMPs-based | 1.119 [1.012–1.238] | 0.029 | 0.874 (0.038) |
|  | DVPs-based | 1.120 [1.010–1.242] | 0.032 | 0.855 (0.038) |
|  | Frailty Index | 1.060 [1.010–1.112] | 0.019 | 0.831 (0.072) |
|  | GrimAge acceleration | 1.034 [0.979–1.091] | 0.233 | 0.829 (0.054) |
|  | Methylation Risk Score | 1.054 [0.937–1.185] | 0.383 | 0.810 (0.063) |
|  | Cognitive function | 0.964 [0.915-1.016] | 0.173 | 0.810 (0.079) |
|  | PhenoAge | 1.021 [0.937-1.113] | 0.635 | 0.801 (0.073) |
|  | Physical function | 0.998 [0.934-1.067] | 0.954 | 0.799 (0.076) |
|  | eFRS | 1.007 [0.520-1.949] | 0.982 | 0.798 (0.074) |
|  | Shannon entropy | 0.973 [0.813–1.163] | 0.761 | 0.798 (0.074) |
|  | Hannum clock | 0.986 [0.870-1.118] | 0.826 | 0.797 (0.074) |
|  | Horvath clock | 0.943 [0.818-1.088] | 0.423 | 0.796 (0.076) |
|  | Null model | – | – | 0.796 (0.075) |

<sup>a</sup> All models are adjusted for age and sex; the null model includes only age and sex covariates (used as predictors). Estimates are based on a sample of 187 individuals with 14 deaths observed over six years.
<sup>b</sup> P-values are from two-sided Wald tests.

We evaluated time-to-event associations between the DMP-DVP epigenetic biomarker and five incident age-related adverse outcomes with at least five events over six years of follow-up: cancer, cardiovascular disease, chronic kidney disease, diabetes, and chronic obstructive pulmonary disease (COPD). After adjustment for age and sex, the biomarker was significantly associated only with an increased hazard of COPD after correction for multiple testing (HR = 1.11, 95% CI: 1.03–1.19, *p* = 0.02, N = 157, 19 events; Table S17). For all outcomes except cancer, models including the biomarker showed higher concordance than the corresponding null models.

The heritability was estimated at 0.41 (SE = 0.37), suggesting that approximately 41% of the variance in epigenetic age deviation is attributable to common SNPs.

Association analysis showed that the DMP-DVP biomarker was positively associated with baseline smoking status (β = 0.0401, *p* = 0.003), high-sensitivity C-reactive protein (β = 0.0471, *p* = 0.032), and interleukin-6 (β = 0.0351, *p* = 0.034) after Bonferroni correction.

### The integrated DMP-DVP biomarker was associated with mortality in BLSA

In external validation using the Baltimore Longitudinal Study of Aging, the integrated DMP-DVP biomarker was associated with all-cause mortality in age- and sex-adjusted Cox models (N = 728; 284 deaths; HR = 1.011, 95% CI: 1.001–1.020, *p* = 0.025; C-index = 0.834). As in CLSA validation set, the integrated biomarker showed higher discrimination than biomarkers constructed from DMPs alone or DVPs alone, supporting the complementary contribution of variability-derived CpGs to mortality-related signal. This pattern was also observed within ancestry-stratified analyses, although overall discrimination was lower in the African ancestry cluster (Table S18).

## Discussion

This study shows that DNA methylation variability may provide a complementary epigenetic signature of aging-related health heterogeneity. In blood samples from older adults, differential methylation and variance association analyses identified largely distinct CpGs, regions, and genes associated with health decline. This pattern is consistent with prior evidence that methylation variability can reflect regulatory instability and that sites showing altered variability are often distinct from those showing shifts in methylation level (3,43). In the present study, this distinction was observed across health deficit accumulation, cognitive function, and physical function, suggesting that variability marks a separate dimension of epigenetic dysregulation in aging.

The biological context differed between differentially methylated and differentially variable regions. Genes overlapped by differentially methylated regions were enriched for immune and inflammatory processes, consistent with immune remodeling in aging-related health decline (44). One of the strongest differentially methylated regions overlapped *SLC1A5*, a glutamine transporter with roles in cellular metabolism and immune-cell function. This is consistent with prior epigenetic frailty work, in which CpGs annotated to *SLC1A5* contributed to a frailty risk score (45). Variability-associated regions did not show broad pathway-level enrichment, but were more localized, CpG-island enriched, and partly shared across health domains. A recurrent variability-associated region overlapped *PSENEN*, a modulator of innate and adaptive immune responses (46), across all three health measures.

Our findings suggest that variability may reflect declining maintenance of epigenetic stability with age. This interpretation is consistent with recent work describing age-related methylation change as quasi-stochastic, where some genomic regions, including promoter CpG islands, may be more susceptible to methylation change, and neighboring CpGs do not necessarily change in a coordinated way (47). The enrichment of variability-associated regions in CpG islands in our study suggests that health-associated methylation instability may be concentrated in regulatory contexts where altered methylation could influence transcriptional regulation (48).

Aggregating these signals into an individual-level biomarker further supported the complementarity of the two classes of CpGs. The combined score showed higher concordance than scores based on differentially methylated or differentially variable CpGs alone. This pattern was also observed in BLSA, although with weaker overall performance, likely reflecting cross-cohort differences and the need to restrict validation to CpGs available on the 450K array.

Several limitations should be considered. The mortality analysis in CLSA was based on a small number of deaths, limiting precision and formal comparison among biomarkers. Cause of death information is unavailable in CLSA, preventing assessment of imbalance between training and validation subsets. Accordingly, mortality associations should be interpreted cautiously, and non significant results for established clocks likely reflect limited power rather than disagreement with prior literature. Although the combined DMP–DVP biomarker showed consistent associations across cohorts with higher concordance, the limited number of deaths reduces statistical precision. These findings therefore require confirmation in larger cohorts with substantially more mortality events. External validation required restriction to CpGs available on the 450K array and involved differences in cohort composition, follow-up duration, and outcome ascertainment. Because analyses were conducted in blood, methylation signals may reflect both cell-intrinsic epigenetic regulation and shifts in immune-cell composition. Finally, the observational design does not establish whether methylation variability contributes to health decline or reflects downstream consequences of aging-related physiological change.

In summary, this study provides supportive evidence that methylation variability may represent a complementary dimension of epigenetic aging. Together, our findings suggest that methylation variability may reflect regionally structured epigenetic dysregulation rather than diffuse genome-wide noise, though larger studies with more adverse events will be needed to assess robustness.

## Acknowledgements

We thank Dr. Andrew Paterson for his thoughtful feedback and productive discussions, which contributed to the development of this manuscript. This research was made possible using the data/biospecimens collected by the Canadian Longitudinal Study on Aging (CLSA). Funding for the Canadian Longitudinal Study on Aging (CLSA) is provided by the Government of Canada through the Canadian Institutes of Health Research (CIHR) under grant reference: LSA 94473 and the Canada Foundation for Innovation, as well as the following provinces, Newfoundland, Nova Scotia, Quebec, Ontario, Manitoba, Alberta, and British Columbia. This research has been conducted using the CLSA Baseline Comprehensive Dataset version 7.0, Follow-up 1 Comprehensive version 4.0, Follow-up 2 Comprehensive version 1.0, Genomic data version 3.0, Epigenetics version 1.1 under Application Number 2206033. The CLSA is led by Drs. Parminder Raina, Christina Wolfson and Susan Kirkland. The time and commitment of the participants to the CLSA study platform is gratefully acknowledged, without whom this research would not be possible.

## Author Contributions

L.T.E. and A.B.W. developed and directed the project. O.V. developed and performed the modeling pipeline and wrote the original draft. A.Z.M. and T.T. performed external validation analyses. J.M., X.S. and K.R. provided critical feedback on the study. O.V. and L.T.E. prepared the manuscript with input from all authors and all authors approved the final manuscript.

## Data availability

Data are available from the Canadian Longitudinal Study on Aging (https://www.clsa-elcv.ca/) for researchers who meet the criteria for access to de-identified CLSA data. Data from the Baltimore Longitudinal Study of Aging are available through submission of research proposals at https://www.blsa.nih.gov/.

## Code availability

The code to analyze the data from both cohorts is available in the GitHub repository https://github.com/oliversion/epg-aging.

## Disclaimer

The opinions expressed in this manuscript are the authors’ own and do not reflect the views of the Canadian Longitudinal Study on Aging. The findings and conclusions presented in this paper are those of the author(s) and do not necessarily reflect the views of the NIH or the U.S. Department of Health and Human Services.

