## Supplementary material for "DNA methylation variability provides a complementary epigenetic signature of aging heterogeneity: Findings from the Canadian Longitudinal Study on Aging and the Baltimore Longitudinal Study of Aging": Figure S1-S5, Supplementary Note 1-2

### Supplementary Notes

**Supplementary Note 1. BLSA study sample and key covariates.** The BLSA analytic sample was constructed from DNA methylation (DNAm) participant-visits with available GWAS ancestry information. Duplicate samples were removed. For participants with more than one eligible DNAm visit, the most recent visit was selected. This DNAm visit was used as the index visit for linkage to other study data and was treated as the time origin for survival analyses. The final study sample included 728 participants with index visits occurring between April 2003 and March 2010.

Mortality status was defined using the last known study status. Participants were coded as deceased if their last known status was deceased. Follow-up time was calculated as the interval between the index visit and death, or between the index visit and last known status for censored participants, using time from study entry as the common time scale. For incident chronic disease outcomes, follow-up time was calculated analogously as the interval from the index visit to the first incident disease visit, or to last known status among participants who remained free of the condition.

Incident dementia was defined as the first observation of adjudicated dementia after the index visit among participants without dementia at the index visit. Adjudicated dementia was identified using `cogn_status = 2`.

Incident type 2 diabetes was identified using a previously defined diabetes variable, `glucose3cat = 2`. Diabetes was defined as present if any of the following criteria were met: fasting glucose $\geq 126$ mg/dL, HbA1c $\geq 6.5\%$, or reported use of diabetes medication.

Incident chronic obstructive pulmonary disease (COPD) was defined as the first observation after the index visit of any of the following: FEV1/PFT $< 0.7$; self-reported COPD during the medical interview together with use of a related medication; or an ICD-9 diagnosis code consistent with COPD, together with use of a related medication. COPD-related ICD-9 codes included 491, 492, and 494.

Incident chronic kidney disease (CKD) was defined as the first observation after the index visit of any of the following: estimated glomerular filtration rate (eGFR) $< 60$ mL/min/1.73 $m^2$, calculated using the CKD-EPI 2021 equation; self-reported CKD during the medical interview; or an ICD-9 diagnosis code consistent with CKD. CKD-related ICD-9 codes included 581, 582, 583, 585, 586, 4030, 4040, 4049, 5900, 5938, 7530, 7531, 7533, and V420.

Incident cardiovascular disease (CVD) was defined as the first observation after the index visit of any of the following: ankle-brachial index (ABI) $< 0.9$; self-reported history during the medical interview of bypass surgery, carotid endarterectomy, aortic aneurysm repair, myocardial infarction, congestive heart failure, angina, stroke, transient ischemic attack, or peripheral artery disease; an ICD-9 diagnosis code consistent with CVD; use of a medication related to congestive heart failure, based on a previously defined medication file indicator (`chfmeds_file = 1`); or use of medications with ATC codes related to CVD. CVD-related ICD-9 codes included 430, 431, 432, 433, 434, 435, 436, 437, 438, 402, 404, 416, 425, 428, 410, 411, 412, 413, 414, 443, 4170, 4291, 4293, V421, V428, V434, 2506, and 4402. CVD-related ATC codes included B01AC23, C01DA, and C01EB18.

**Supplementary Note 2. CpG-level overlap between DMP-DVP biomarker and established epigenetic clocks.** We investigated the overlap between DNA methylation sites used as predictors in epigenetic biomarkers. After variable selection using elastic net regression, 789 distinct CpG sites were retained in the model for the DVP-DMP-based epigenetic biomarker. Given the regional correlation structure of DNA methylation, exact overlap in CpG sites between clocks is not necessarily expected. Consistent with this, only 8 CpG sites were shared between our proposed biomarker and the established GrimAge and PhenoAge clocks. To assess potential functional convergence, we examined gene annotations for the CpGs included in each clock. This analysis identified only two genes, *HDAC4* and *SMPD3*, that were common to all three clocks.

### Supplementary Figures

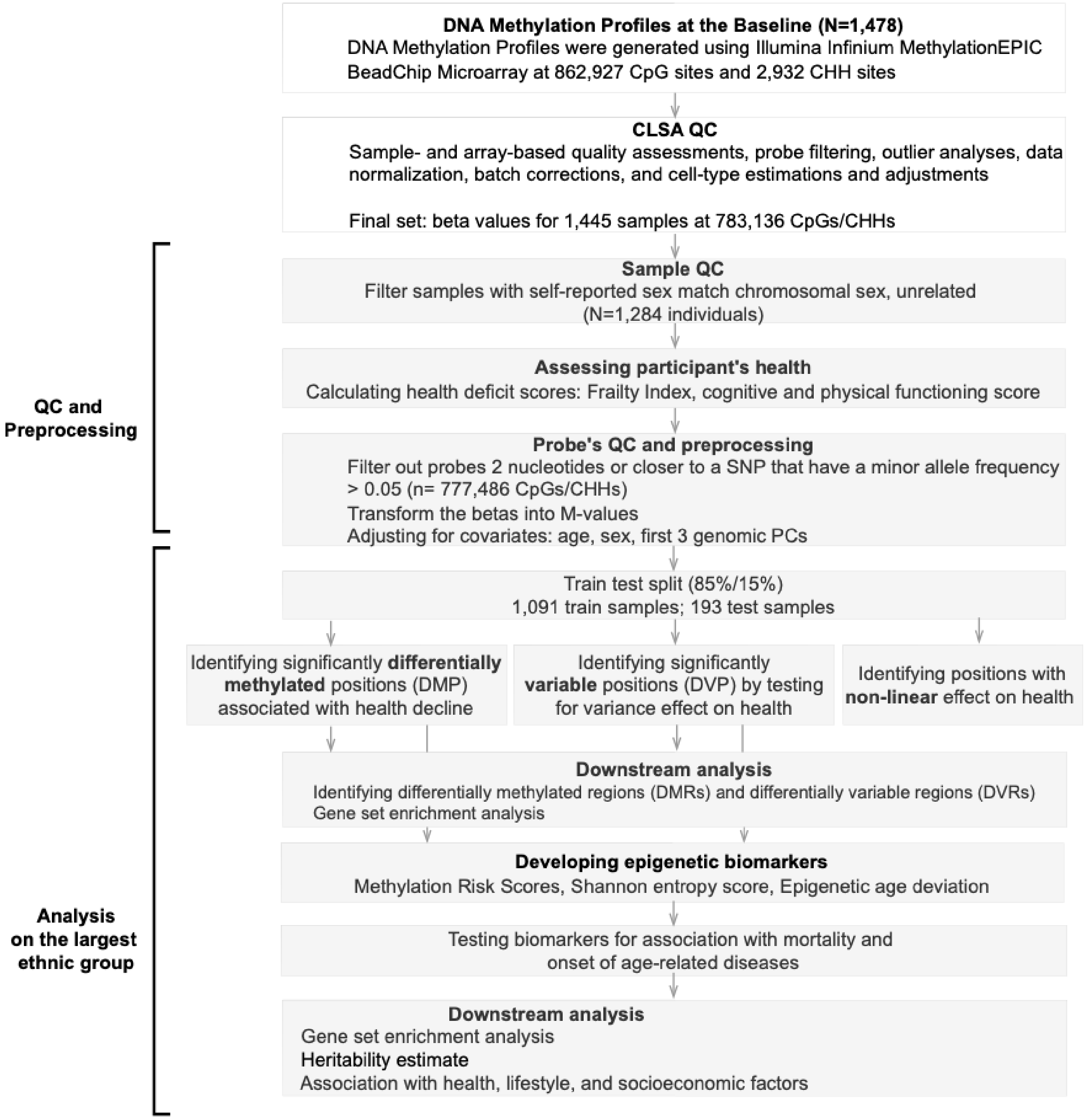


**Figure S1.** Study workflow.

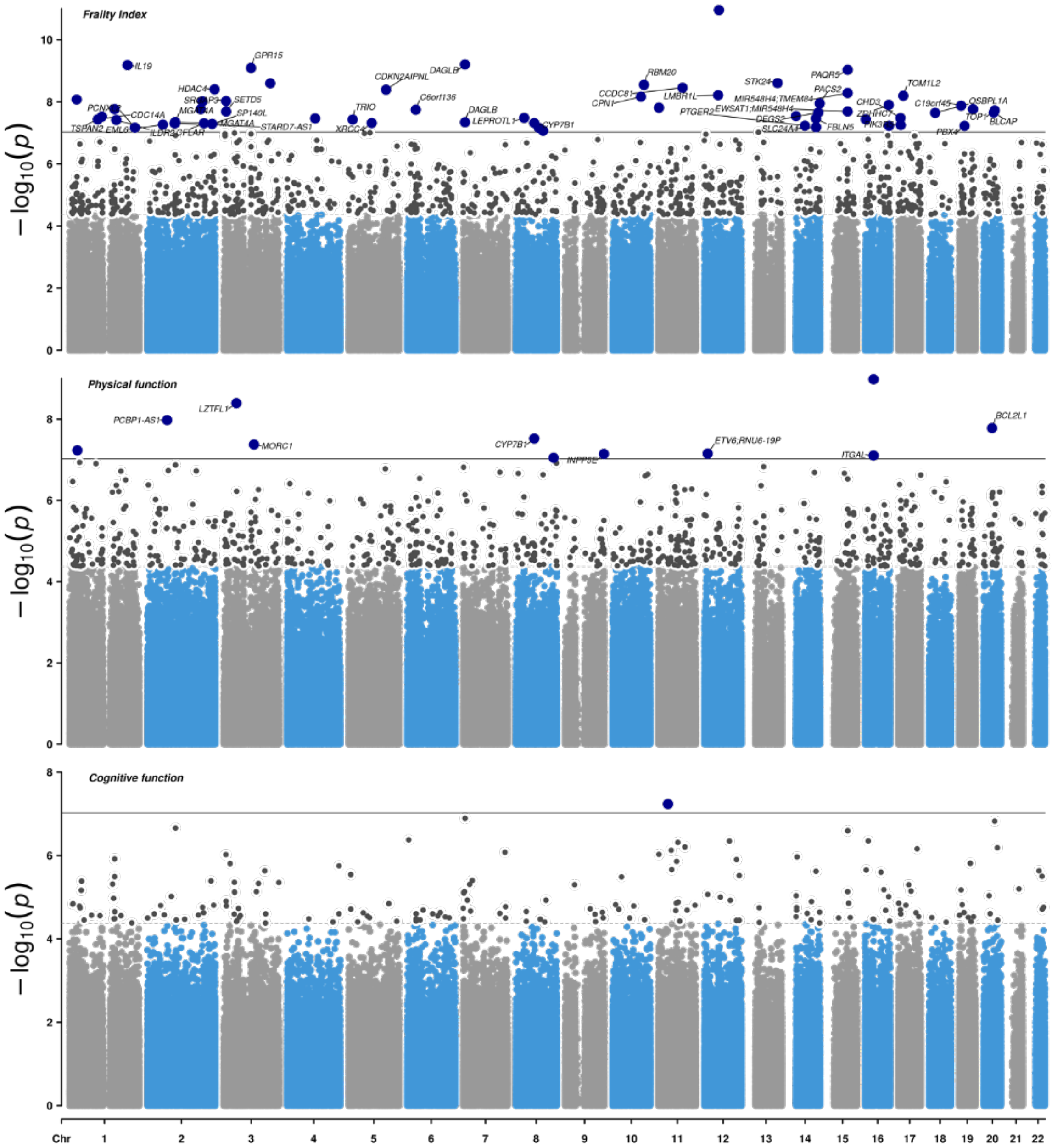


**Figure S2.** Genome-wide differential methylation analysis across three continuous health deficit scores identified 1,775 differentially methylated positions (DMPs) that exceeded FDR correction, of which 70 surpassed the epigenome-wide significance threshold ($9.42 \times 10^{-8}$). P-values are from a two-sided z-test. The black line denotes the epigenome-wide significance threshold. The gray dashed line denotes the FDR threshold.

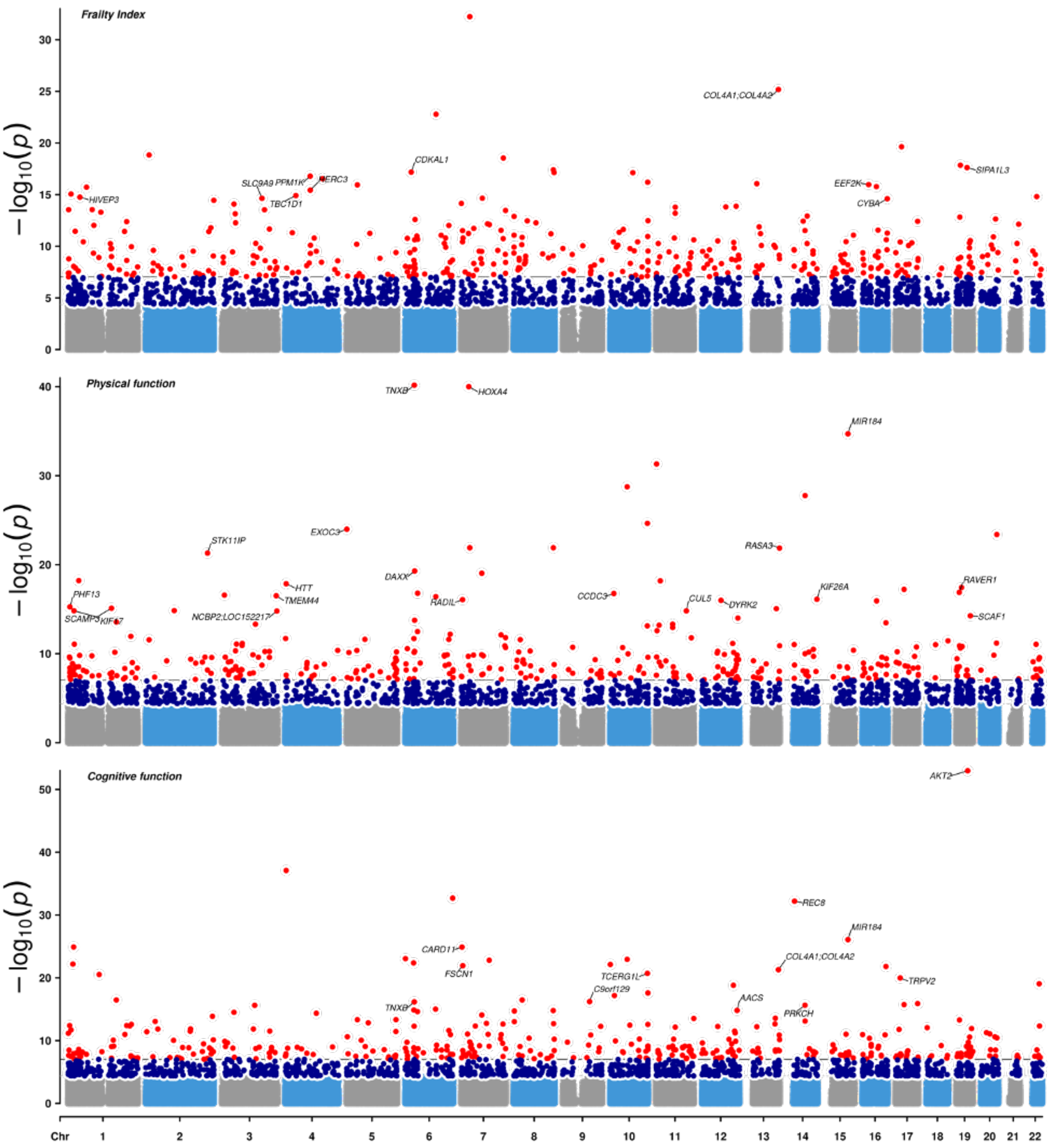


**Figure S3.** Genome-wide variance association analysis for DNA methylation sites across three continuous health deficit scores identified 3,197 differentially methylated positions (DVPs) that exceeded FDR correction, of which 473 surpassed the epigenome-wide significance threshold ($9.42 \times 10^{-8}$). $-\log_{10}(p$-values) are from a two-sided z-test derived from the dispersion model of double generalized linear models (DGLMs), testing the association between methylation variance (M-values) and each health deficit score. The blue dashed line denotes the epigenome-wide significance threshold ($p < 9.42 \times 10^{-8}$). The gray dashed line denotes the FDR threshold.

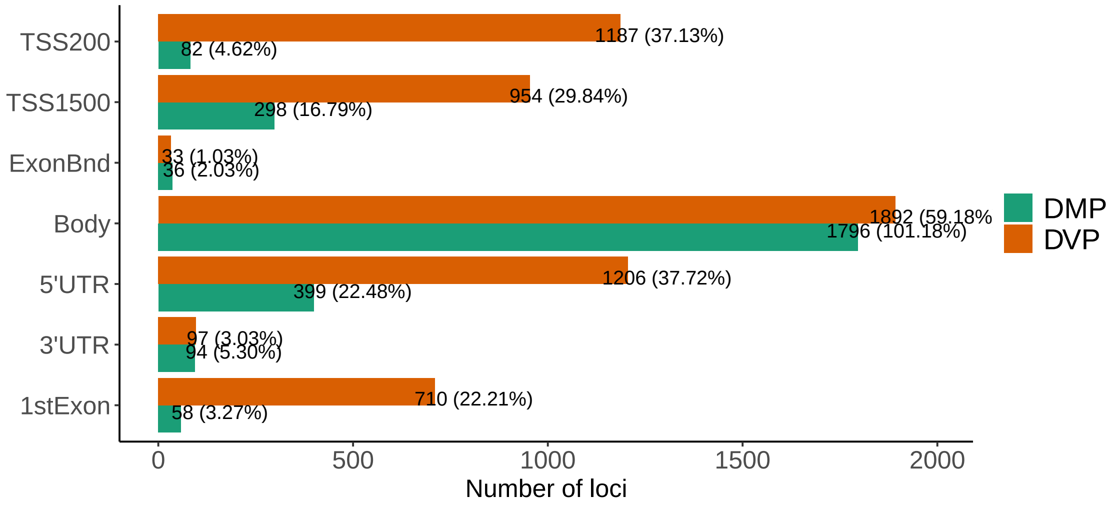


**Figure S4.** Functional genomic region annotation are shown for differentially methylated positions (DMPs), differentially variable positions (DVPs), and all tested CpGs from the Illumina EPIC v1 array background. The background included 764,839 CpGs. Percentages were calculated within each CpG set: 1,775 DMPs, 3,197 DVPs, and 764,839 tested EPIC v1 CpGs. CpG annotations were grouped as Island, Shore, Shelf, and OpenSea.

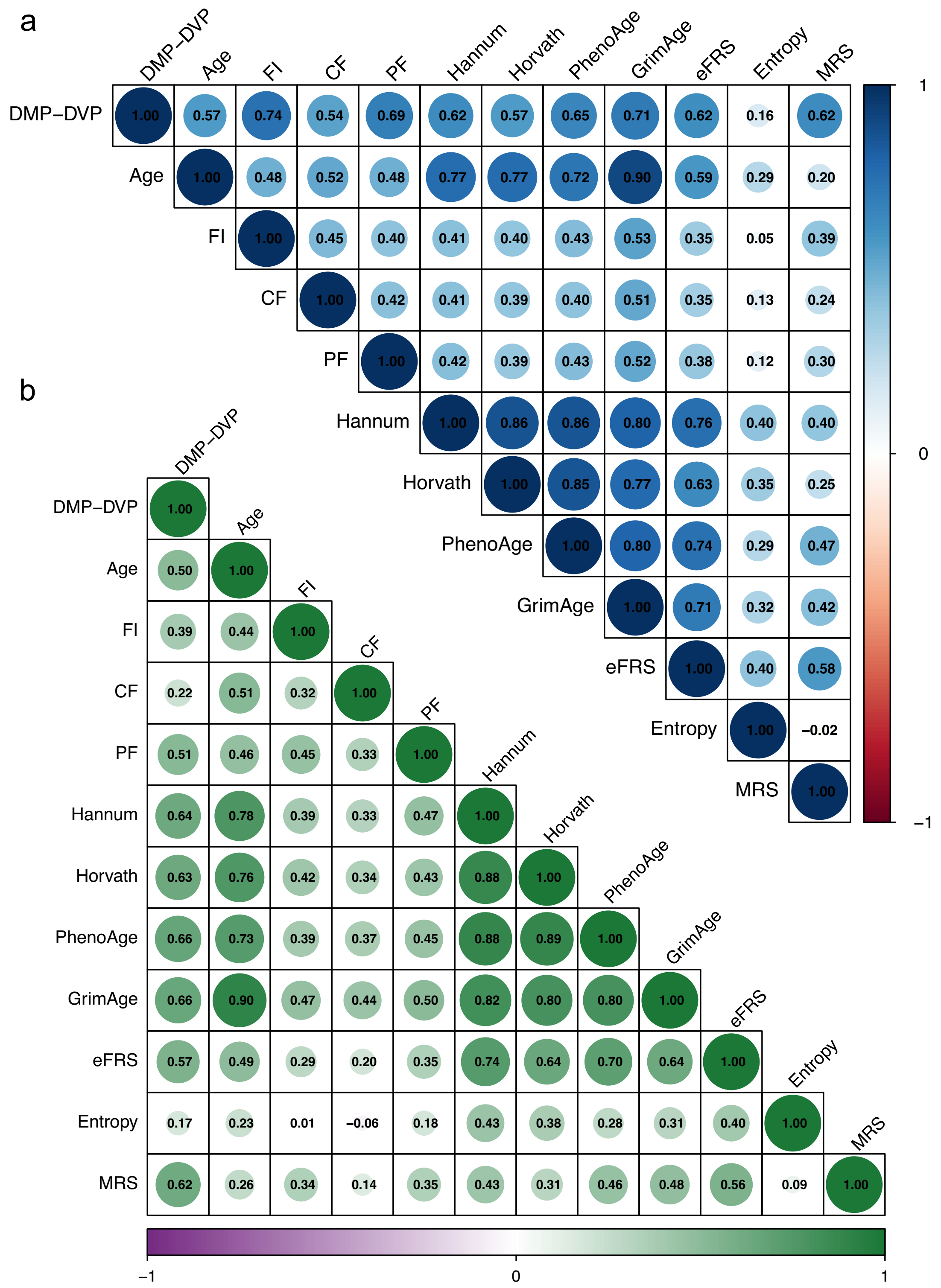


**Figure S5.** Pairwise Pearson correlations between mortality predictors in both the CLSA training and validation sets. DMP-DVP epigenetic score includes CpGs identified from both differential methylation and differential variability analysese. A methylation Shannon entropy measure calculated over DVPs. FRS - the epigenetic frailty risk score; MRS - the methylation risk score; FI - Frailty Index; CF - cognitive function; PF - physical function
